# Late sporogonic stages of *Plasmodium* parasites are susceptible to the melanization response in *Anopheles gambiae* mosquitoes

**DOI:** 10.1101/2024.05.31.596773

**Authors:** Suheir Zeineddine, Sana Jaber, Sally A. Saab, Johnny Nakhleh, George Dimopoulos, Mike A. Osta

**Author notes:** **Correspondence:** Mike A. Osta. These authors contributed equally to this work (Suheir Zeineddine, Sana Jaber).

## Abstract

The malaria-causing parasites have to complete a complex infection cycle in the mosquito vector that also involves attack by the insect’s innate immune system, especially at the early stages of midgut infection. However, *Anopheles* immunity to the late *Plasmodium* sporogonic stages, such as oocysts, has received little attention as they are considered to be concealed from immune factors due to their location under the midgut basal lamina and for harboring an elaborate cell wall comprising an external layer derived from the basal lamina that confers self-properties to an otherwise foreign structure. Here, we investigated whether *Plasmodium berghei* oocysts and sporozoites are susceptible to melanization-based immunity in *Anopheles gambiae*. Silencing of the negative regulator of melanization response, CLIPA14, increased melanization prevalence without significantly increasing the numbers of melanized oocysts, while co-silencing CLIPA14 with CLIPA2, a second negative regulator of melanization, resulted in a significant increase in melanized oocysts and melanization prevalence. Only late-stage oocysts were found to be melanized, suggesting that oocyst rupture was a prerequisite for melanization-based immune attack, presumably due to the loss of the immune-evasive features of their wall. We also found melanized sporozoites inside oocysts and in the hemocoel, suggesting that sporozoites at different maturation stages are susceptible to melanization. Silencing the melanization promoting factors TEP1 and CLIPA28 rescued oocyst melanization in CLIPA2/CLIPA14 co-silenced mosquitoes. Interestingly, silencing of CTL4, that protects early stage ookinetes from melanization, had no effect on oocysts and sporozoites, indicating differential regulation of immunity to early and late sporogonic stages. Similar to previous studies addressing ookinete stage melanization, the melanization of *Plasmodium falciparum* oocysts was significantly lower than that observed for *P. berghei*. In summary, our results provide conclusive evidence that late sporogonic malaria parasite stages are susceptible to melanization, and we reveal distinct regulatory mechanisms for ookinete and oocyst melanization.

## 1 Introduction

*Anopheles* mosquitoes are not passive vectors of malaria parasites, but rather parasites suffer significant losses in numbers during all developmental stages, which explains why several mosquitoes remain uninfected in highly endemic regions (Gouagna et al., 2010; Mbogo et al., 1993; Mendis et al., 2000). The transformation of ookinetes to oocysts in the basal labyrinth of the midgut epithelium constitutes the most severe bottleneck during *Plasmodium* development, often reducing the numbers of developing oocysts to single digits (Gouagna et al., 1998; Sinden, 1999). Survival of ookinetes is determined by a fine balance between antagonistic and agonistic immune factors that regulate two key effector responses against *Plasmodium* ookinetes in the midgut tissue: melanization and lysis.

The complement-like protein TEP1 is, perhaps, the most studied antagonist described to date that is responsible for the killing of the majority of invading ookinetes as they contact the hemolymph in the basal labyrinth of the midgut epithelium (Blandin et al., 2004). The active form of TEP1 circulates in the hemolymph as a complex with the two leucine-rich proteins LRIM1 and APL1C that are important to stabilize TEP1 in circulation (Fraiture et al., 2009; Povelones et al., 2009). TEP1 binds to ookinetes of the rodent malaria parasite *Plasmodium berghei* triggering their lysis, and its knockdown (kd) in susceptible and resistant strains of *An. gambiae* results in a significant increase in oocyst numbers (Blandin et al., 2004; Blandin et al., 2009). TEP1 activation to mediate its anti-*Plasmodium* effects seems to depend also on microvesicle release by midgut-homing hemocytes through an unknown mechanism (Castillo et al., 2017). Although TEP1 is also involved in the killing of ookinetes of the human malaria parasite *P. falciparum* (Dong et al., 2006), it seems that different *P. falciparum* strains exhibit different levels of susceptibility to TEP1 through mechanisms that are not completely understood (Eldering et al., 2016; Molina-Cruz et al., 2012; Molina-Cruz et al., 2013). TEP1 is also essential for *P. berghei* and *P. falciparum* ookinete melanization, however, its contribution seems to depend on the refractory mosquito genetic background; it is essential for melanizing ookinetes of both species in L3-5 mosquitoes (Collins et al., 1986; Blandin et al., 2004; Eldering et al., 2016), however, it is dispensable for *P. falciparum* ookinete melanization in CTL^null^ mosquitoes (Simoes et al., 2022). The clip domain serine proteases (cSPs) and their serine protease homologs (cSPHs) that are central regulators of the melanization response constitute another class of *Plasmodium* antagonists, as several members are critical for *Plasmodium* melanization (El Moussawi et al., 2019; Povelones et al., 2013; Sousa et al., 2020; Volz et al., 2006; Zhang et al., 2021), in particular the three cSPHs SPCLIP1, CLIPA8 and CLIPA28 which form a hierarchical cSPH module that is central for the activation of the protease cascades driving microbial melanization (El Moussawi et al., 2019; Saab et al., 2024; Sousa et al., 2020).

The major *Plasmodium* agonists include the C-type lectins CTL4 and CTLMA2 that function together as a complex (henceforth CTL complex) (Osta et al., 2004; Schnitger et al., 2009; Simoes et al., 2022; Simoes et al., 2017) and the non-catalytic clip domain serine protease homologs (cSPHs) CLIPA2 (Volz et al., 2006; Yassine et al., 2014) and CLIPA14 (Nakhleh et al., 2017). These agonists function as negative regulators of the mosquito melanization response, and their silencing triggers a potent melanotic response against *P. berghei* ookinetes residing in the basal labyrinth with a concomitant reduction in the numbers of live oocysts (Nakhleh et al., 2017; Osta et al., 2004; Volz et al., 2006; Yassine et al., 2014). *CTL4* kd mosquitoes trigger significant *P. falciparum* melanization only at high infection levels (Simoes et al., 2017), whereas CTL4^null^ transgenic mosquitoes, in which the CTL4 gene is disrupted, melanized all *P. berghei* ookinetes (100% refractoriness) and a substantial number of *P. falciparum* ookinetes even at low infection intensities that mimic those reported in field caught mosquitoes, reducing infection prevalence (i.e. % of midguts with at least 1 live oocyst) 2-3 folds (Simoes et al., 2022). These results indicate that melanization is so far the most potent anti-*Plasmodium* defense system in the mosquito that may be exploited to hamper malaria transmission. While TEP1 is essential for *P. berghei* ookinete melanization in *CTL4* kd mosquitoes (Povelones et al., 2011), the melanization of *P. falciparum* ookinetes in CTL4^null^ mosquitoes is TEP1-independent (Simoes et al., 2022), indicating that the response to these two *Plasmodium* species is differentially regulated. CLIPA2 and A14 synergistically regulate *P. berghei* ookinete melanization, as co-silencing both genes trigger a potent melanotic response to *P. berghei* reducing infection prevalence of the mosquito to 26%, compared to 86% in controls (Nakhleh et al., 2017). However, it remains unknown whether *CLIPA2*/*A14* double kd (dkd) mosquitoes also melanize *P. falciparum* ookinetes.

Studies of mosquito immune responses to *Plasmodium* have largely focused on ookinetes being the most vulnerable stage. Immune responses to oocysts, however, remain largely unknown despite the fact that oocysts mature during a period of 10 days providing ample time for interactions with both the humoral and cellular components of the host immune response. *Plasmodium* oocysts exhibit reductions in numbers between day 2 (early oocyst development) and day 8 in *An. gambiae* mosquitoes and this has been partially attributed to functions mediated by STAT and Litaf-like 3 (LL3) transcription factors; although the exact mechanisms remain to be characterized, LL3 and STAT seem to independently control hemocyte differentiation in response to parasite infection (Gupta et al., 2009; Smith et al., 2015). A recent study revealed that *P. berghei* infection in *An. stephensi* drives the JAK/STAT pathway-dependent proliferation of midgut enteroblasts that surround oocysts triggering their elimination by lysis and phagocytosis (Barletta et al., 2024). Here, we aimed to characterize the susceptibility of *Plasmodium* late sporogonic stages, mature oocysts and sporozoites, to melanization in *An. gambiae CTL4* kd and *CLIPA2*/*CLIPA14* dkd mosquitoes, which are known to exhibit potent melanotic refractoriness to ookinete stages (Osta et al., 2004; Simoes et al., 2022; Simoes et al., 2017; Nakhleh et al., 2017). We show that mature oocysts and sporozoites of wildtype *P. berghei* are indeed susceptible to the melanization response, but contrary to our expectations, CTL4, CLIPA2 and CLIPA14 appear to exhibit *Plasmodium* stage-specific roles in the context of the melanization response. Only *CLIPA2*/*CLIPA14* dkd mosquitoes melanized a significant number of *P. berghei*, and to a lesser extent, *P. falciparum* oocysts. Oocyst melanization occurred at a late stage concomitant with oocyst wall rupture, as young oocysts observed at day 10 pi (pi) did not exhibit significant melanization.

## 2 Materials and Methods

### 2.1 Rearing of *Anopheles gambiae* mosquitoes

Adult female mosquitoes of the *Anopheles gambiae* G3 were used for all *P. berghei* infections and Keele strain was used for *P. falciparum* infections. Mosquitoes were maintained in the insectary at 27°C, 80% humidity, with a 12-hour day-night cycle. Larvae were contained in plastic pans and fed with Tetra Pond flakes/sticks. Emerging adults were collected from the larval pans into small cages and fed on a 10% sucrose solution. For egg laying, female mosquitoes were fed on BALB/c mice anesthetized by a mixture of ketamine/xylazine.

### 2.2 Mosquito infections with *Plasmodium* and scoring of parasites numbers

Mosquito infections with *P. berghei* were done using GFP-expressing *P. berghei* parasite (PbGFPCON) (Franke-Fayard et al., 2004) by allowing batches of wild type mosquitoes to feed on anesthetized, 5 to 6-week-old BALB/c mouse infected with *P. berghei* at 4%-6% blood parasitemia for approximately 10 minutes at 20°C. Mosquitoes were thereafter kept at 20°C until dissected. At day 7 post-infection (p.i.) with *P. berghei*, mosquitoes were injected with the corresponding dsRNA for gene silencing, and midguts were dissected at days 10 and 14 to assess melanization in intact and ruptured oocysts, respectively. To assess the effect of an additional blood meal on the intensity of oocyst melanization at day 14 p.i., the same infection protocol above was followed except that mosquitoes were given an additional naïve blood meal at day 9 p.i. All midguts were fixed for 30 min in 4% paraformaldehyde, washed 3 times with 1x phosphate-buffered saline (PBS) then mounted in Fluoroshield histology mounting medium (Sigma). For each experiment, 2 to 4 independent biological repeats were performed. The statistical difference in the numbers of oocysts between the different groups was performed using Kruskal-Wallis test followed by Dunn’s multiple comparisons test, with *P*-values less than 0.05 considered significant. Statistical analysis for melanization prevalence (i.e. % of midguts with at least 1 melanized oocyst) was determined by Fisher’s exact test with *P*-values less than 0.05 considered significant. Raw data of parasite counts are shown in Supplementary Table 1.

To count *P. berghei* salivary gland sporozoites, heads were gently detached from the mosquito thoraces at day 21 p.i. ensuring that all 6 lobes remain attached. Each head was grinded separately in 50 μl of 1x sterile PBS using a motor grinder to release sporozoites from glands. The corresponding homogenates were centrifuged at 100g for 4 minutes at 4°C. Supernatants were collected into fresh tubes, and 10 μl of each sample were loaded into a hemocytometer to count the numbers of GFP-expressing sporozoites using a Leica upright fluorescent microscope. The statistical difference in the numbers of sporozoites between the different groups was performed using Kruskal-Wallis test followed by Dunn’s multiple comparisons test, with *P*-values less than 0.05 considered significant. To acquire images of melanized sporozoites attached to salivary glands, heads with glands attached were dissected from mosquito thoraces at day 21 p.i. as described above, fixed and washed as described for the midguts. However, before mounting, glands were separated from heads and images were acquired on a Leica upright fluorescent microscope.

Infections with *P. falciparum* were performed by feeding ds*GFP* (control group) and ds*CLIPA2*/*A14 An. gambiae* (Keele strain) female mosquitoes on *P falciparum* NF54 gametocyte culture with human blood at 27°C through artificial glass feeders two days post-dsRNA injection. Different dilutions of gametocytemia were used to achieve different infection intensities. To be able to achieve a high infection intensity with a high oocyst median, the mosquitoes were fed on antibiotic-treated 10 % sucrose containing 25 μg/ml gentamicin sulfate and 100 units-μg/ml of penicillin–streptomycin throughout the infection assay. Mosquito guts were then dissected seven days post-infection and the counts of both live and melanized parasites were scored as previously described (Simoes et al., 2022). For late-stage melanization experiments, adult female mosquitoes were first fed on *P falciparum*-infected blood meal, then injected with the appropriate dsRNA five days p.i. and the guts were dissected 12 days p.i. Statistical analysis of mean parasite numbers was done using the non-parametric Mann-Whitney test with *P-values* less than 0.05 were considered significant. Statistical analysis for prevalence was tested using Fisher’s exact test with *P-values* less than 0.05 were considered significant.

### 2.3 Synthesis of double-stranded RNA and gene silencing by RNAi

Synthesis of the double-stranded RNAs (dsRNAs) used to silence the genes of interest was done using the T7 RiboMax Express Large-Scale RNA production system (Promega) according to the instructions of the manufacturer. dsRNAs were purified using phenol:chloroform:isoamyl alcohol (25:24:1), followed by precipitation by isopropanol. dsRNA pellets were then resuspended in nuclease free water, and their concentration was finally adjusted to 3.5 μg/μL. Primers used in dsRNA synthesis are listed in Supplementary Table 2.

*In vivo* gene silencing was performed as previously described (Blandin et al., 2002). Briefly, 1 to 3-days-old female adult mosquitoes were anesthetized over CO_2_, and microinjected intrathoracically with 69 nL of the corresponding gene-specific dsRNA at 3.5 μg/μL concentration for single knock downs, or with 138 nL of a 1:1 mixture of two different dsRNAs each at 3.5 μg/μL concentration for double knock downs. For triple gene knock downs, equal volumes of three different dsRNAs at 4.5 μg/μL concentration each, were mixed and 138 nL from the corresponding mixture were injected. Injections were performed using the Drummond Nanoject II Nanoliter injector.

### 2.4 Determination of gene silencing efficiency

The efficiency of gene silencing was assessed at the protein level by Western Blot analysis. Hemolymph was extracted into 1x Laemmli buffer (Bio-Rad) by proboscis clipping at day 7 post dsRNA injection. Hemolymph protein samples from around 20 females were resolved by SDS-PAGE, followed by wet transfer to immuno-blot PVDF membrane (Bio-Rad). Membranes were blocked with 1x phosphate-buffered saline (PBS) containing 0.05% Tween 20 and 3% milk solution for 1 hour at room temperature, followed by an overnight incubation at 4°C with the corresponding primary antibodies. Primary antibodies were used according to the following dilutions: rabbit αTEP1, 1:1000 (El Moussawi et al., 2019), rabbit αCTL4, 1:1000 (Saab et al., 2024), rabbit αCLIPA14, 1:3000 (Saab et al., 2024), rabbit αCLIPA2, 1:1000 (Yassine et al., 2014), rabbit αCLIPA28, 1:1000 (El Moussawi et al., 2019), and mouse αApolipophorin II, 1:100 (Mendes et al., 2008). Next, membranes were washed with 1xPBS-Tween 20 and incubated with secondary anti-rabbit (1:14,000) or anti-mouse (1:6000) antibodies conjugated with horse-radish peroxidase for 1 hour at room temperature. Membranes were revealed with Bio-Rad Clarity Max Western ECL substrate and imaged using the ChemiDoc MP System (Bio-Rad).

## 3 Results

### 3.1 Mature oocysts and sporozoites of wild-type *P. berghei* are susceptible to the mosquito melanization response

The melanization of *P. berghei and P. falciparum* ookinetes and early oocysts has been extensively studied, however, to what extent the late stages of malaria parasites are susceptible to the mosquito melanization response has remained unknown. To address this question, female *An. gambiae* G3 strain mosquitoes were first given an infectious blood meal containing *P. berghei*, then, 7 days later, they were injected with dsRNAs (ds) specific to the key negative regulators of ookinete melanization including, CLIPA2 (Yassine et al., 2014), CLIPA14 (Nakhleh et al., 2017) and CTL4 (Osta et al., 2004; Simoes et al., 2022; Simoes et al., 2017), in order to trigger the melanization response against late oocysts. Efficient silencing of all 3 genes was confirmed by western blot analysis (Supplementary Figure 1). Melanized, mature oocysts were scored in midguts dissected at day 14 post-*P. berghei* infection, the time at which most oocysts exhibit different stages of wall rupture, according to a detailed scanning electron microscopy study of *P. berghei* oocyst development in *An. gambiae* (Orfano et al., 2016). The mean numbers of melanized oocysts per midgut in ds*CLIPA2* and ds*CLIPA14* mosquitoes were not significantly different from the background level in ds*LacZ* controls (Figure 1A, scatter plot). However, melanization prevalence (i.e. percentage of mosquitoes carrying at least 1 melanized oocyst) was significantly higher in ds*CLIPA14* (40%) compared to ds*LacZ* (9%) mosquitoes (Figure 1A, pie charts), suggesting that *CLIPA14* kd does enhance the melanization of oocysts, yet not dramatically. Interestingly, co-silencing both genes triggered a significant increase in the mean number of melanized oocysts at day 14 pi (henceforth day-14 oocysts) and in melanization prevalence (Figures 1A, C). Ds*CLIPA2*/*A14* (i.e. received a mixture of dsRNAs specific to CLIPA2 and CLIPA14) mosquitoes exhibited a mean number of live oocysts that was systematically lower than that in ds*LacZ* control, however the difference was not statistically significant possibly due to the large variation in the efficiency of oocyst melanization between individual midguts. Only in rare cases (10% of midguts) ds*CLIPA2*/*A14* mosquitoes exhibited total melanotic refractoriness. Contrary to our expectations, ds*CTL4* mosquitoes exhibited background levels of oocyst melanization (Figures 1B, C), indicating that CTL4 is not involved in regulating oocyst melanization, in contrast to its pivotal regulatory role in ookinete melanization (Osta et al., 2004; Simoes et al., 2022; Simoes et al., 2017). This result is also in agreement with a lack of oocyst melanization in CTL4^null^ *A. gambiae* (Simoes et al., 2022).

**Figure 1.**
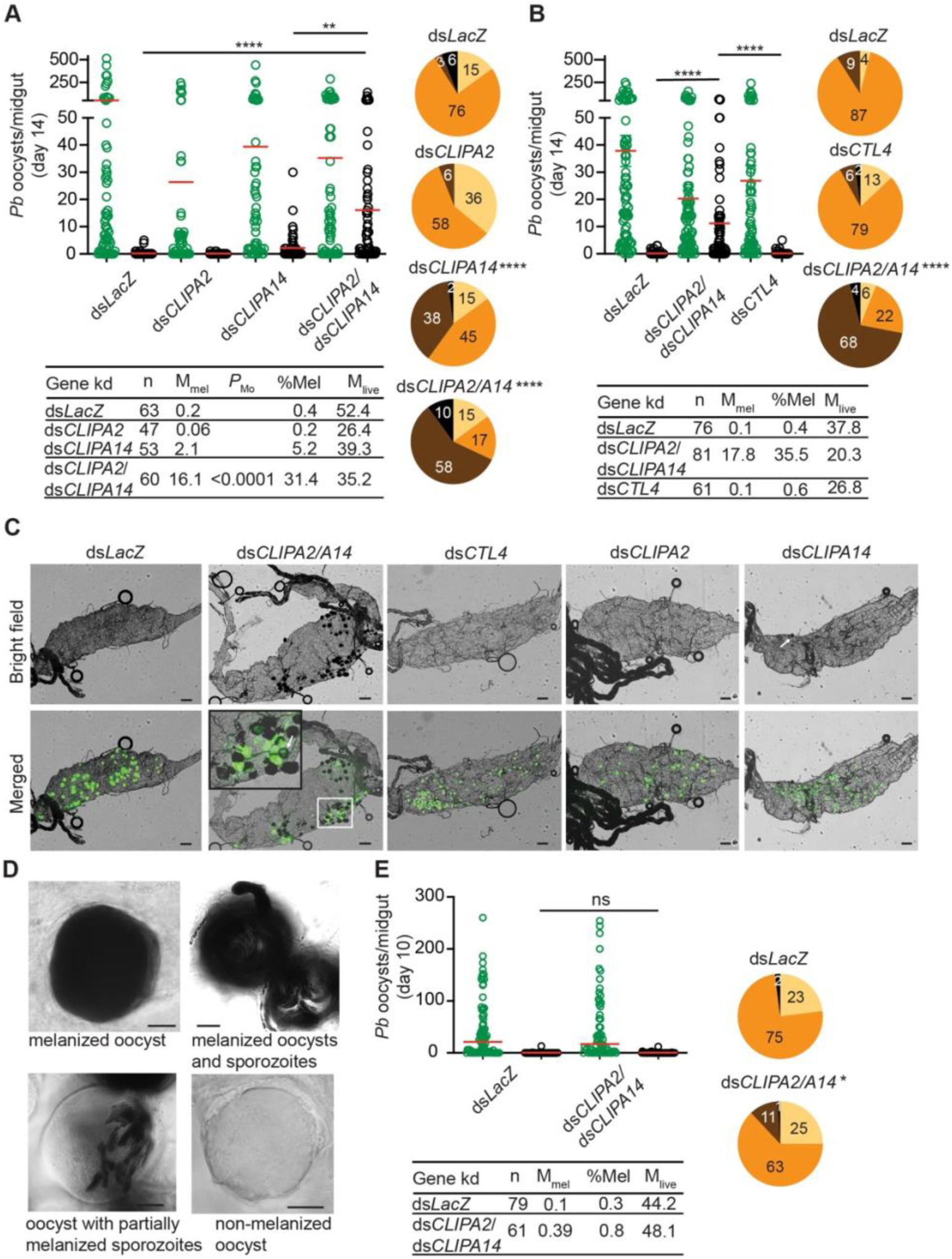
Co-silencing CLIPA2 and CLIPA14 trigger the melanization of *P. berghei* late oocysts and sporozoites. (**A** and **B**) Mosquitoes were injected with the indicated dsRNAs at day 7 after receiving an infectious blood meal and midguts were dissected at day 14 post-blood feeding to score the numbers of live (GFP-expressing; green circles) and melanized (black circles) *P. berghei* oocysts. Red lines on the scatter plots indicate mean parasite numbers. Statistical analysis was performed using Kruskal-Wallis test followed by Dunn’s multiple comparisons test, with *P*-values less than 0.05 considered significant. The tabulated data under each figure show the percentage of melanized oocysts (% Mel) and the mean numbers of melanized (M_mel_) and live (M_live_) oocysts per midgut. *n*, number of midguts analyzed. Pie charts show the percentages of midguts that are non-infected (beige), infected with live oocysts only (orange), carrying both live and melanized parasites (dark brown) or only melanized parasites (black) in each gene kd; *P*-values for pie charts were determined by Fisher’s exact test using as one of the categorical variables the presence or absence of melanized oocysts. Data shown are from 4 and 2 independent trials for **(A)** and **(B)**, respectively. **(C)** Representative microscopy images of whole midguts of the indicated mosquito genotypes constructed by tiling. Shown are bright field and merged (bright field with green channel) images. The area defined by a white rectangle in dsCLIPA2\A14 image is shown as an enlarged inset. White arrows point to melanized oocysts. Scales bars, 100 μm. **(D)** Representative images of day-14 oocysts in ds*CLIPA2*/*A14* mosquitoes with different levels of melanization. Scale bars, 10 μm. **(E)** Numbers of live (green circles) and melanized (black circles) *P. berghei* oocysts in the midguts of ds*CLIPA2*/*A14* mosquitoes dissected at day 10 after infection. Data shown are from 3 independent experiments. Statistical analysis, labelling of the accompanying tabulated data and pie charts are as in **A** and **B**. ****, *P*<0.0001; **, *P*<0.01; *, *P*<0.05; ns, non-significant.

A more careful observation of melanized, day-14 oocysts at high magnification revealed that the sporozoites inside them are also melanized (Figure 1D). Oocysts exhibited various degrees of melanization, with some completely melanized while others showing partial melanization (Figure 1D). To determine whether oocyst susceptibility to melanization requires prior rupturing of the oocyst wall, we assessed the melanization of oocysts at day 10 pi (henceforth day-10 oocysts) in ds*CLIPA2*/*A14* mosquitoes, using the same experimental approach described above. Indeed, the mean number of melanized oocysts in these mosquitoes was similar to control, and melanization prevalence was marginally significant (Figure 1E) and much lower than that scored for day-14 oocysts (compare ds*CLIPA2*/*A14* pie charts in Figure 1E with those in Figures 1A-B), suggesting that oocysts’ susceptibility to the melanization response is dramatically enhanced after their walls start to rupture. This also suggests that melanization of mature, ruptured oocysts is initiated on the parasite-derived inner wall layers, either on sporozoite surfaces or on the inner sporoblast membranes (Thathy et al., 2002), but not on the mosquito-derived outer wall layer. The image of the oocyst with partially melanized sporozoites in Figure 1D supports this conclusion. As noted in this figure, no melanization is detected around the oocyst, yet several sporozoites appear melanized inside it. Also, in the inset of the representative ds*CLIPA2*/*A14* midgut image in Figure 1C, the arrow points to a partially melanized oocyst with melanin deposits all around its rim, yet GFP is still detected in its center; had melanization started on the outer surface of this oocyst prior to rupturing, the melanin coat would have masked the GFP signal emitted from inside the oocyst.

Melanization in ds*CLIPA2*/*A14* mosquitoes does not only target sporozoites in the context of ruptured oocysts but also circulating sporozoites. Indeed, melanized sporozoites were observed attached to the salivary glands of ds*CLIPA2*/*A14* but not ds*LacZ* or ds*CTL4* mosquitoes at day 21 pi with *P. berghei* (Figure 2B). At day 21 pi, melanized sporozoites were also detected in the abdomens of ds*CLIPA2*/*A14* mosquitoes (Figure 2C), and oocyst and sporozoite melanization was extensive in the midguts of these mosquitoes at that late time point (Figure 2D). Consequently, ds*CLIPA2*/*A14* mosquitoes harbored significantly fewer salivary gland sporozoites at day 21 compared to the ds*LacZ* and ds*CTL4* groups (Figure 2A). Altogether, these results indicate that sporozoites are targeted by melanization from the early stage of egress from oocysts until they reach the salivary glands.

**Figure 2.**
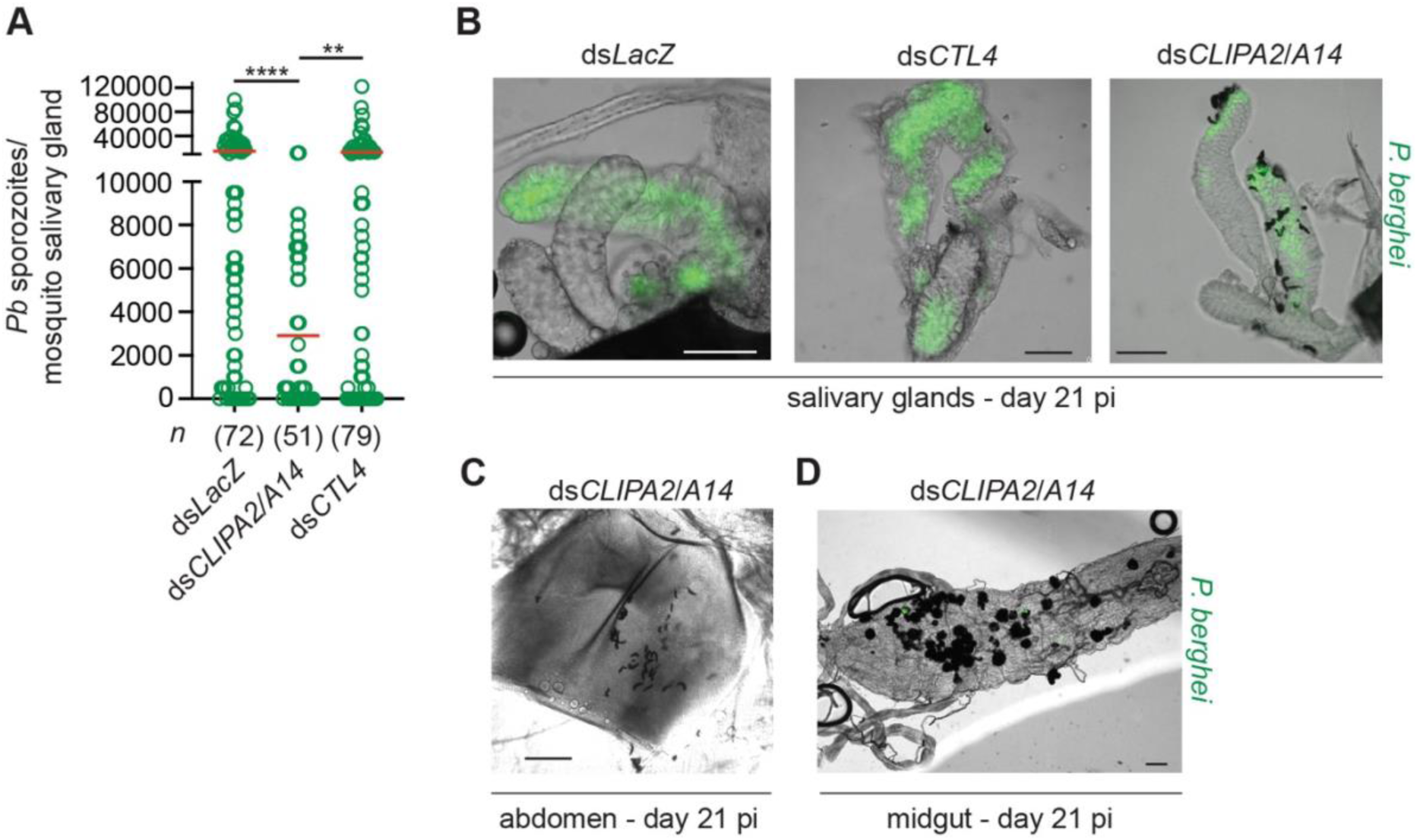
Ds*CLIPA2*/*A14* mosquitoes harbor fewer salivary gland sporozoites. **(A)** Scatter plot showing the numbers of *P. berghei* sporozoites scored in salivary glands dissected at day 21 from the indicated mosquito genotypes. Red lines on the scatter plots indicate mean parasite numbers. Data shown are from 4 independent biological experiments. Statistical analysis was performed using Kruskal-Wallis test followed by Dunn’s multiple comparisons test, with *P*-values less than 0.05 considered significant. ****, *P*<0.0001; **, *P*<0.01. *n*, numbers of salivary glands counted per sample. **(B)** Representative images showing melanized sporozoites attached to the salivary glands at day 21 pi with *P. berghei* (green) in ds*CLIPA2*/*A14* but not in ds*LacZ* or ds*CTL4* mosquitoes. **(C-D)** Representative images showing (C) Melanized sporozoites in a ds*CLIPA2*/*A14* abdomen, and **(D)** melanized oocysts from a ds*CLIPA2*/*A14* midgut, dissected at day 21 pi with *P. berghei*. Scale bars, 75 μm.

Mosquitoes take multiple blood meals in the field. Since blood feeding was shown to increase the numbers of circulating hemocytes (Baton et al., 2009; Bryant and Michel, 2014; Castillo et al., 2006) which are an important source of several factors of melanization including, clip domain serine proteases, TEP1, and PPOs among others (Kwon et al., 2021; Pinto et al., 2009; Raddi et al., 2020), we hypothesized that an additional blood meal might enhance the melanization of mature oocysts in ds*CLIPA2*/*A14* mosquitoes. However, giving mosquitoes an additional naive blood meal 3 days post-dsRNA injection (i.e. at day 10 after the first infectious blood meal) did not enhance the melanization of day-14 oocysts (Figure 3), suggesting that the partial melanization phenotype in these mosquitoes is not due to a shortage of immune factors driving melanization.

**Figure 3.**
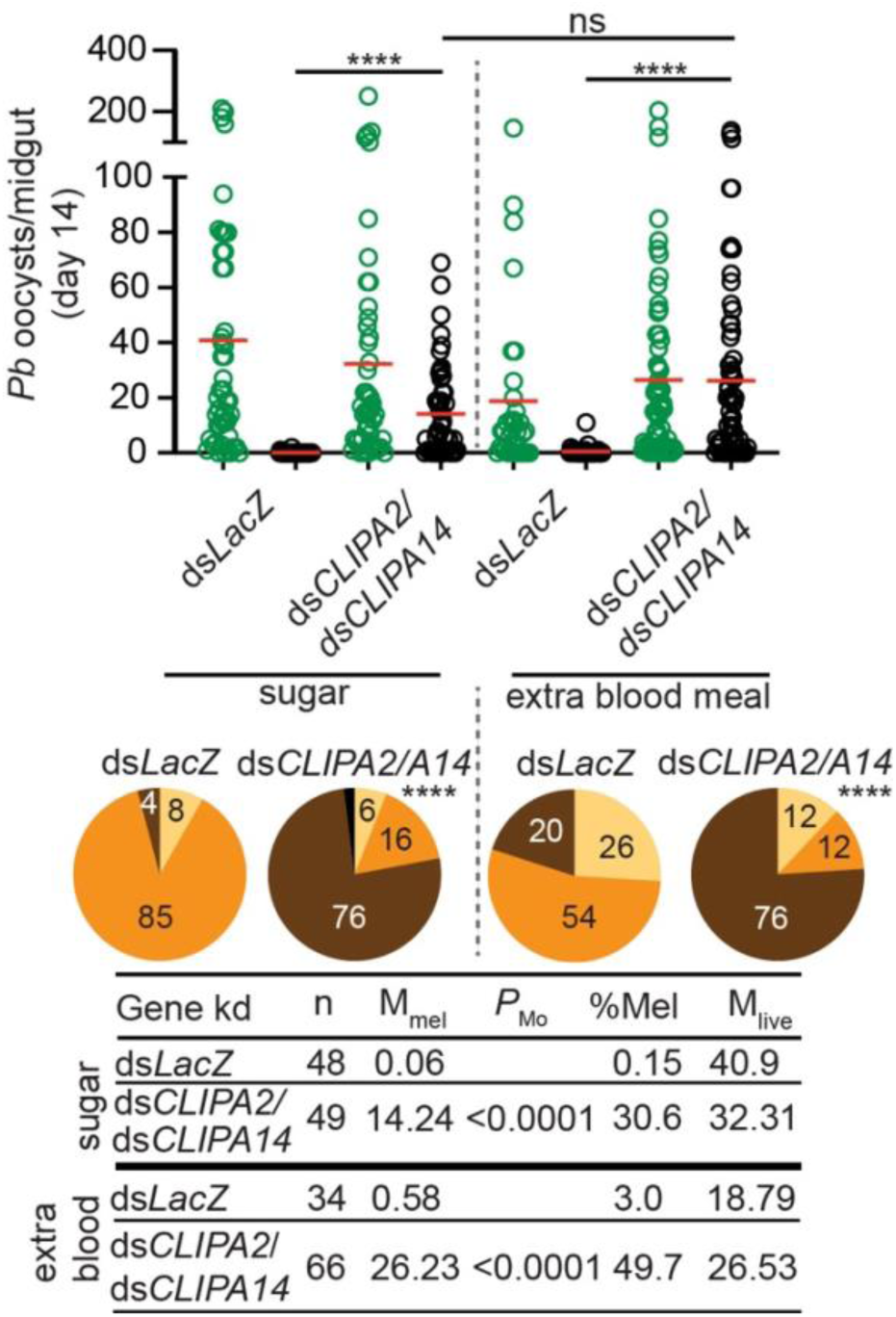
An additional bloodmeal does not enhance oocyst melanization in ds*CLIPA2*/*A14* mosquitoes. Mosquitoes were injected with ds*CLIPA2*/*A14* at day 7 after receiving *P. berghei*-infected blood. One group was kept on sugar and the second was given an additional naïve blood meal at day 10 after the first infectious blood meal (i.e. 3 days after dsRNA injection). Midguts were dissected at day 14 post-*P. berghei* infection to score the numbers of live (GFP-expressing; green circles) and melanized (black circles) *P. berghei* oocysts. Red lines on the scatter plots indicate mean parasite numbers. Statistical analysis was performed using Kruskal-Wallis test followed by Dunn’s multiple comparisons test, with *P*-values less than 0.05 considered significant. The tabulated data show the percentage of melanized oocysts (% Mel) and the mean numbers of melanized (M_mel_) and live (M_live_) oocysts per midgut. *n*, number of midguts analyzed. Pie charts show the percentages of midguts that are non-infected (beige), infected with live oocysts only (orange), carrying both live and melanized parasites (dark brown) or only melanized parasites (black) in each gene kd; *P*-values for pie charts were determined by Fisher’s exact test using as one of the categorical variables the presence or absence of melanized oocysts. Data shown are from 3 independent trials. ****, *P*<0.0001.

### 3.2 *P. berghei* oocyst melanization in ds*CLIPA2*/*A14* mosquitoes requires TEP1 and the cSPH module

The observed *CTL4* RNAi phenotype suggests that the melanization of mature oocysts and ookinetes may be subject to distinct regulatory mechanisms. To further investigate this point, we checked whether oocyst melanization in ds*CLIPA2*/*A14* mosquitoes can be reversed after silencing TEP1 and CLIPA28, both of which are essential factors in *An. gambiae* melanization response to diverse microbes (Blandin et al., 2004; Eldering et al., 2016; El Moussawi et al., 2019; Nakhleh et al., 2017; Povelones et al., 2011; Sousa et al., 2020; Yassine et al., 2012). Indeed, the mean number of melanized oocysts in ds*CLIPA2*/*A14* mosquitoes was dramatically reduced after silencing TEP1 or CLIPA28 (Figures 4A-B); melanization prevalence in ds*CLIPA2*/*A14*/*A28* mosquitoes was similar to ds*LacZ*, while that in ds*CLIPA2*/*A14*/*TEP1* mosquitoes remained slightly higher than ds*LacZ*, yet significantly lower than ds*CLIPA2*/*A14* (Figure 4, compare pie charts). These results indicate that TEP1 and CLIPA28 are essential for oocyst melanization in ds*CLIPA2*/*A14* mosquitoes. No increase in the numbers of live oocysts was noted in ds*CLIPA2*/*A14*/*TEP1* mosquitoes which is in line with a previous study showing that TEP1 is not involved in the late phase immune response to developing oocysts (Smith et al., 2015).

**Figure 4.**
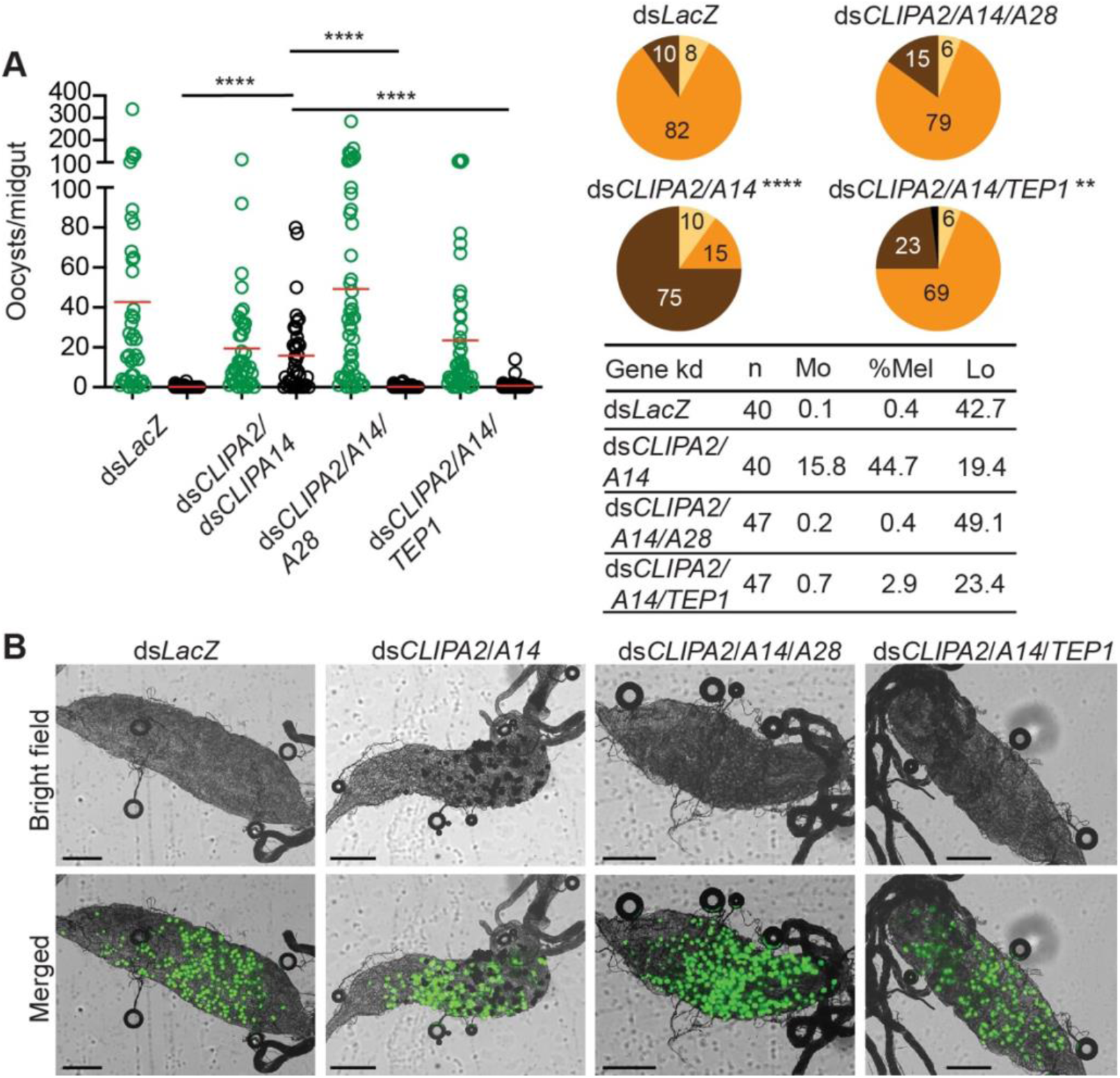
TEP1 and CLIPA28 are required for oocyst melanization in mosquitoes co-silenced for CLIPA2 and CLIPA14. **(A)** Mosquitoes were injected with the indicated dsRNAs at day 7 after receiving an infectious blood meal and midguts were dissected at day 14 post-blood feeding to score the numbers of live (GFP-expressing; green circles) and melanized (black circles) *P. berghei* oocysts. Red lines on the scatter plots indicate mean parasite numbers. Statistical analysis was performed using Kruskal-Wallis test followed by Dunn’s multiple comparisons test, with *P*-values less than 0.05 considered significant. The tabulated data show the percentage of melanized oocysts (% Mel) and the mean numbers of melanized (M_mel_) and live (M_live_) oocysts per midgut. *n*, number of midguts analyzed. Pie charts show the percentages of midguts that are non-infected (beige), infected with live oocysts only (orange), carrying both live and melanized parasites (dark brown) or only melanized parasites (black) in each gene kd; *P*-values for pie charts were determined by Fisher’s exact test using as one of the categorical variables the presence or absence of melanized oocysts. Data shown are from 2 independent trials. **(B)** Representative microscopy images of whole midguts of the indicated mosquito genotypes constructed by tiling. Shown are bright field and merged (bright field with green channel) images. Scales bars are 250 μm. ****, *P*<0.0001; **, *P*<0.01.

### 3.3 *P. falciparum* ookinetes and mature oocysts are susceptible to the melanization response in ds*CLIPA2*/*A14* mosquitoes

*P. falciparum* ookinetes are susceptible to the melanization response triggered in *CTL4*-silenced (Simoes et al., 2017) and CTL4^null^ mosquitoes (Simoes et al., 2022), however, to lower levels than those of *P. berghei*. To address the effect of co-silencing *CLIPA2* and *CLIPA14* on *P. falciparum* infection, we first addressed the impact on the early-stage ookinetes in *An. gambiae* Keele strain mosquitoes. Adult female mosquitoes injected with ds*CLIPA2*/*A14* or ds*GFP* (control group) were infected with the *P. falciparum* NF54 strain through feeding on a gametocyte culture and their midguts were dissected seven days post-infection to assess infection phenotypes. Co-silencing of CLIPA2 and CLIPA14 triggered significant melanization of early *P. falciparum* stages (Figure 5A, left panel, and Supplementary Table 3) with no effect on the infection prevalence (Figure 5A, middle panel). Melanized parasites were detected in the midguts of 5% of ds*CLIPA2*/*A14* mosquitoes (Figure 5A, right panel). Since *P. falciparum* melanization in Keele *An. gambiae* strain was previously shown to occur more prominently at high-intensity infections (Simoes et al., 2017), we repeated this experiment with a higher infection intensity. The results show that the number of melanized parasites was relatively higher (Figure 5B, left panel) and that the melanized parasites were observed in 15% of the ds*CLIPA2*/*A14* mosquito midguts (Figure 5B, right panel). However, the number of melanized parasites per midgut was still rather low compared to *P. berghei* infections, and both melanized ookinetes and early oocysts were observed (Figure 6, filled and normal arrows, respectively). This phenotype agrees with previous studies (Simoes et al., 2017) showing that melanization is mainly triggered upon higher-intensity *P. falciparum* infections. We then investigated the effect of co-silencing CLIPA2 and CLIPA14 on the mature oocyst stage. To do so, mosquitoes previously infected with *P. falciparum-*infected blood meal were co-injected with ds*CLIPA2* and ds*CLIPA14*, and both live and melanized mature oocysts were counted twelve days post-infection. Our results showed a significant melanization of mature *P. falciparum* oocysts in ds*CLIPA2*/*A14* mosquitoes with a melanization prevalence of 39 % compared to 0% in ds*LacZ* (Figures 5C, 6, arrowhead).

**Figure 5.**
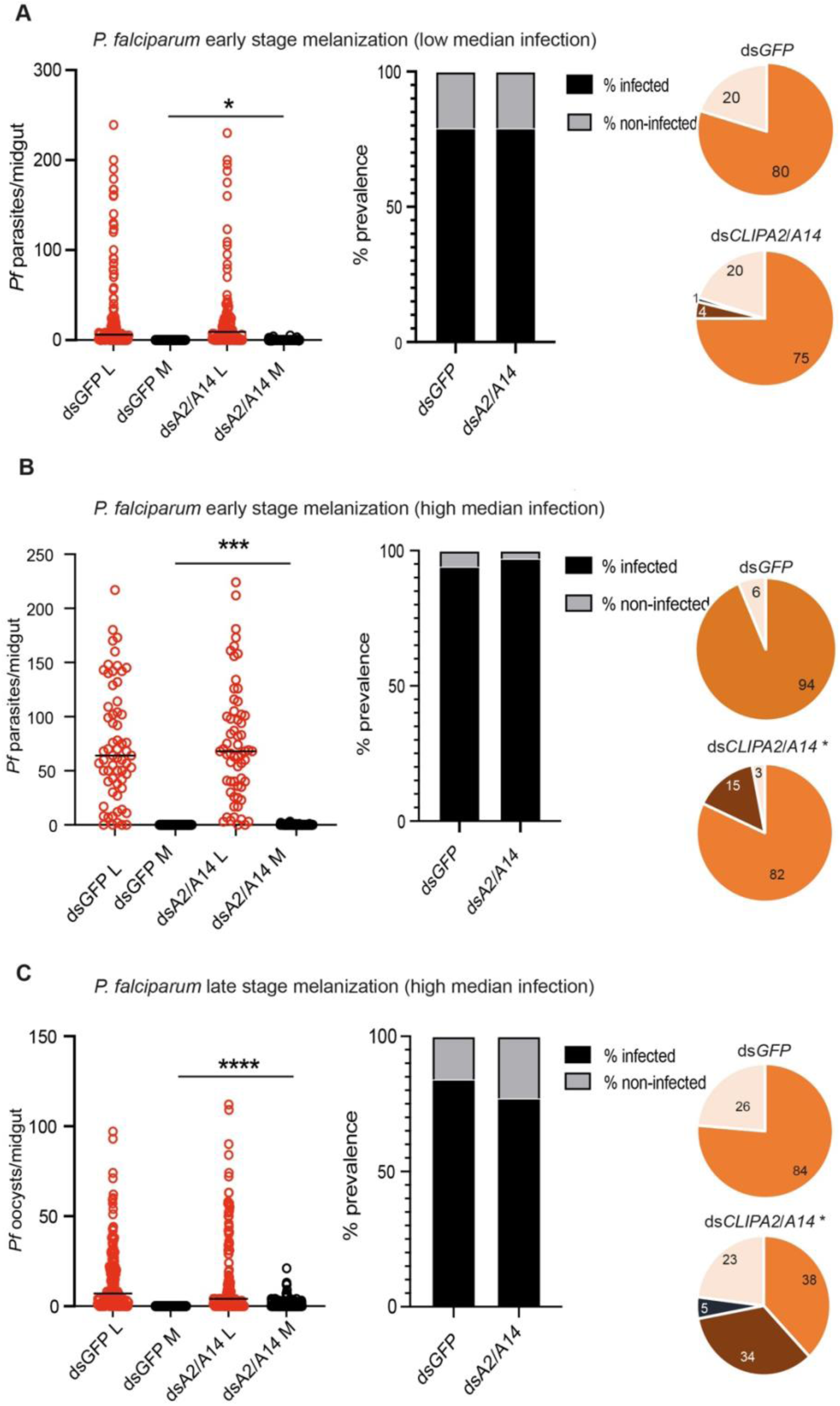
Co-silencing CLIPA2 and CLIPA14 trigger the melanization of both early and late *P. falciparum* midgut stages. Scatter plots (left panels) showing the number of parasites counted in the midguts of the indicated *An. gambiae* mosquito genotypes (Keele strain) (**A** and **B**) seven days PI (low and high median infection respectively) or **(C)** twelve days PI. L, live parasites (red circles), and M, melanized parasites (black circles). Medians were shown as red lines. Statistical analysis was done using the non-parametric Mann-Whitney test, with *P-values* less than 0.05 considered significant. The prevalence bar graphs (middle panel) show the percentage of mosquitoes that are infected with at least one parasite in their guts. Pie charts (last panel) show the percentages of midguts that are non-infected (beige), infected with live oocysts only (orange), carrying both live and melanized parasites (dark brown) or only melanized parasites (black) with an asterisk resembling significance. Statistical analysis for prevalence was tested using Fisher’s exact test with *P-values* less than 0.05 considered significant. *, *P*<0.05; ***, *P*<0.001; ****, *P*<0.0001. *Pf*, *P. falciparum*.

**Figure 6.**
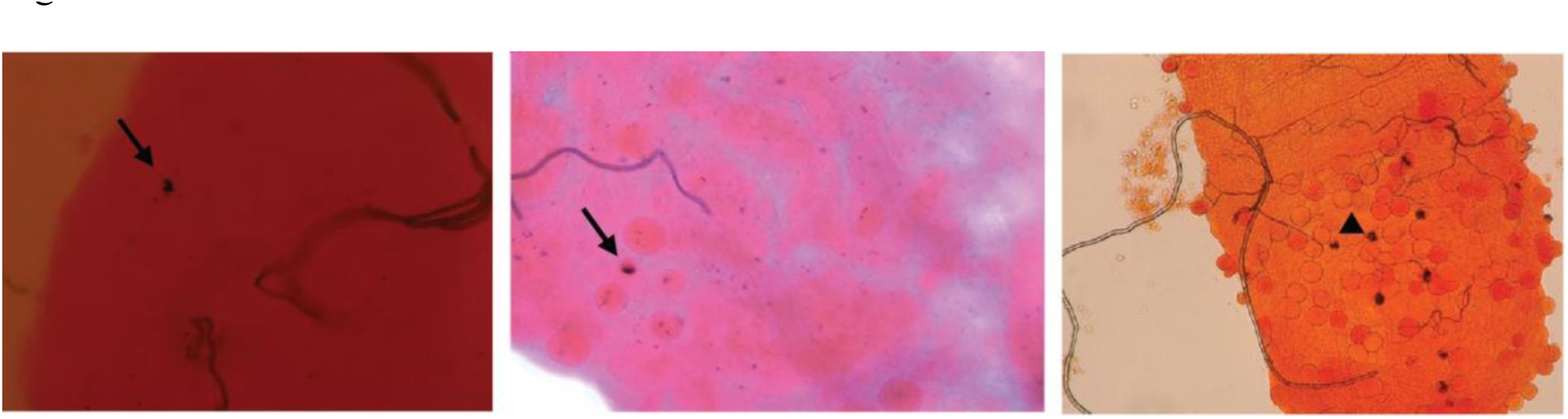
CLIPA2/CLIPA14 double kd mosquito guts showing melanized ookinete (filled arrow), early oocyst (normal arrow), and late oocyst (arrowhead)

## 4 Discussion

The melanization response is likely the most potent anti-*Plasmodium* defense system in the mosquito, with detrimental effects on *Plasmodium* ookinete stage parasites (Nakhleh et al., 2017; Osta et al., 2004; Simoes et al., 2022; Simoes et al., 2017; Yassine et al., 2014), thereby exerting a strong selective pressure on the parasite that may explain why it is rarely observed in nature (Niare et al., 2002; Riehle et al., 2006; Schwartz and Koella, 2002). Here, we extend the characterization of this system to late stages in the parasite’s sporogonic cycle, specifically mature oocysts and sporozoites that have been generally considered more resilient to mosquito immune responses compared to ookinetes. We show that co-silencing CLIPA2 and CLIPA14 triggers significant melanization of *P. berghei* mature oocysts and sporozoites and reduces significantly the numbers of sporozoites that successfully reach and invade the salivary glands. whereas, surprisingly, CTL4 does not seem to regulate the melanization of these late stages, in contrast to its central role in controlling ookinete melanization (Osta et al., 2004; Simoes et al., 2022; Simoes et al., 2017). This differential role of CTL4 in ookinete and oocyst melanization could be due to differences in the genetic interactions that regulate the immune response to these two stages. TEP1 binds *P. berghei* ookinetes triggering their lysis (Blandin et al., 2004) and is also required for *P. berghei* ookinete melanization in *CTL4* kd mosquitoes (Povelones et al., 2011), hence, CTL4 may function as a molecular switch in the TEP1 module to suppress ookinete melanization skewing the response towards lysis. Since TEP1 does not target oocysts (Blandin et al., 2004; Smith et al., 2015), the function of CTL4 as a molecular switch may become dispensable at later stages, explaining plausibly the lack of an RNAi phenotype. Another indication that distinct genetic interactions could be operating at the ookinete and oocyst stages to regulate melanization are the RNAi phenotypes of CLIPA2 and CLIPA14. Silencing either of these genes triggers a dramatic increase in the numbers of melanized *P. berghei* ookinetes, with *CLIPA14* kd exhibiting the most prominent phenotype, resembling that of CTL4 (Nakhleh et al., 2017; Yassine et al., 2014). Co-silencing both genes further enhances *P. berghei* ookinete melanization reducing infection prevalence to 26%, compared to 86% in controls, suggesting that CLIPA2 and CLIPA14 synergistically regulate ookinete melanization (Nakhleh et al., 2017). In this study, the individual knockdowns of these genes did not significantly increase the mean numbers of melanized oocysts per midgut relative to the control group, but only their co-silencing did. These results indicate that CLIPA2 and CLIPA14 exhibit almost complete functional redundancy with respect to oocysts melanization, but unique functions in the context of ookinete melanization. The molecular basis for these phenotypes remains unclear since the mechanisms of action of these cSPHs have not been characterized.

TEP1 does not bind oocysts nor target them for lysis (Blandin et al., 2004; Smith et al., 2015), yet it is still required for their melanization, at least in the context of *P. berghei* infections. Hence, the fact that *TEP1* kd significantly reversed the melanization of oocysts in ds*CLIPA2*/*A14* mosquitoes indicates that TEP1 function is firmly entrenched in the melanization response, supporting previous observations of its indispensable role in bacterial (El Moussawi et al., 2019; Kamareddine et al., 2016; Sousa et al., 2020), fungal (Yassine et al., 2012) and ookinete melanization (Blandin et al., 2004; Nakhleh et al., 2017; Povelones et al., 2011). The only time TEP1 deviated from that role is in CTL4^null^ mosquitoes, whereby it was shown to be dispensable to *P. falciparum* ookinete melanization (Simoes et al., 2022), possibly because *P. falciparum* is more apt at evading the mosquito immune response (Molina-Cruz et al., 2015; Molina-Cruz et al., 2013). This also explains why *P. falciparum* ookinete and oocyst melanization was much weaker than that observed for *P. berghei* (this study and (Nakhleh et al., 2017)). The weaker melanization of *P. falciparum* can be also attributed to the differences in infection temperatures of each parasite species. Mosquito infections with *P. berghei* and *P. falciparum* are carried at 20°C and 27°C degrees, respectively, and infected mosquitoes are maintained at these respective temperatures until dissection, and this results in different infection kinetics that may enable one parasite species to be more efficiently targeted by the melanization system. Accordingly, it was previously shown that the proportion of melanized *P. falciparum* ookinetes in CTL4^null^ mosquitoes was higher at 20°C relative to 27°C degrees (Simoes et al., 2022). Importantly, *A. gambiae* is not a natural mosquito vector species for *P. berghei* as it is for *P. falciparum* that has, therefore, likely developed more sophisticated immune-evasive mechanisms through co-evolution. The fact that CLIPA28 strongly reversed oocyst melanization in ds*CLIPA2*/*A14* mosquitoes lends further support to the central role of the core cSPH module (SPCLIP1-CLIPA8-CLIPA28), in which CLIPA28 is the most downstream member, in the mosquito melanization response to diverse microbial challenges (El Moussawi et al., 2019; Kolli et al., 2022; Povelones et al., 2013; Schnitger et al., 2007; Sousa et al., 2020; Yassine et al., 2012).

The mean numbers of melanized *P. falciparum* and *P. berghei* oocysts in this study, though significant, are considered moderate compared to those reported previously for *P. falciparum* ookinetes in CTL4^null^ mosquitoes (Simoes et al., 2022) and for *P. berghei* ookinetes in CTL4^null^ (Simoes et al., 2022), ds*CTL4* (Osta et al., 2004; Simoes et al., 2017) and ds*CLIPA2*/*A14* mosquitoes (Nakhleh et al., 2017). There are several plausible explanations for this observation that are non-mutually exclusive. First, we have shown here that the melanization of day-10 oocysts was insignificant, suggesting that the rupturing of the oocyst wall precedes melanization. Our results agree with those of a recent study reporting that *P. berghei* and *P. falciparum* oocysts lacking the enzyme glutaminyl cyclase become susceptible to the melanization response only after their walls rupture, whereas no melanization was observed in oocysts before day 10 post-infection (Kolli et al., 2022). Thus, the asynchronous maturation of *Plasmodium* oocysts in the midgut and the fact that not all oocysts would have ruptured at day 14 (Beier, 1998; Vaughan et al., 1994; Zollner et al., 2006; Orfano et al., 2016), some may never rupture at all, could explain the different susceptibilities of individual oocysts to melanization. Second, in addition to asynchronous development, asynchronous gene expression programs at the single oocyst and sporozoite levels may also influence susceptibility to melanization by modulating the interactions with the host. In support of this argument, the glutaminyl cyclase-mutant *P. berghei* and *P. falciparum* parasites in the study of Kolli *et al*. were observed at most in 65% and 34% of infected *An. stephensi* midguts, respectively, and even in those midguts that contained melanized parasites, many were not melanized (Kolli et al., 2022), indicating that several mutant oocysts and sporozoites escaped melanization. These results suggest that susceptibility to melanization cannot be explained by a single parasite trait and differences in gene expression programs between individual oocysts and sporozoites may explain this observation. Indeed, a study employing single-cell RNA sequencing identified extensive transcription heterogeneity among *P. berghei* sporozoites isolated from the same anatomical site (Bogale et al., 2021). Third, the incomplete depletion of CLIPA2 and CLIPA14 by RNAi may also partially explain the moderate oocyst melanization phenotype in ds*CLIPA2*/*A14* mosquitoes, which is one of the drawbacks of RNAi. This is especially critical in the context of CLIPA14, which in our hands is one of the least efficiently silenced hemolymph proteins, as substantial amounts are still detected in the hemolymph by western blot after RNAi ((Nakhleh et al., 2017) and Supplementary Figure 1). The complete knock out of these two genes is expected to reveal more potent phenotypes of *Plasmodium* melanization. Indeed, this was observed with CTL4 whose silencing by RNAi, though highly efficient (Schnitger et al., 2009), triggered strong yet partial melanization of *P. berghei* ookinetes (Osta et al., 2004) and weak melanization of *P. falciparum* ookinetes observed only at high infection levels (Simoes et al., 2017), however, *CTL4* knockout completely melanized *P. berghei* and strongly melanized *P. falciparum* ookinetes (Simoes et al., 2022).

Oocyst wall rupture appears to be a prerequisite to signal parasite melanization in ds*CLIPA2*/*A14* mosquitoes. The oocyst wall is composed of an internal layer of parasite origin and an external layer or capsule derived from the mosquito midgut basal lamina (Aikawa, 1971; Arrighi and Hurd, 2002). According to this and two previous studies utilizing mutant *Plasmodium* parasites (Kolli et al., 2022; Zhu et al., 2022), melanization seems to initiate on the internal oocyst layer and sporozoite membranes rather than on the external layer which probably has, yet to be identified, regulatory proteins that protects it from melanization, like other tissues of the mosquito. The most intriguing question is the nature of the ligand(s) that the melanization response senses on membranes of parasite origin or is it the fact that these membranes intrinsically lack specific host regulatory factors renders them vulnerable to that response. Circumsporozoite (CS) protein, the most abundant protein on sporozoites seems to be a prime candidate in protecting parasites from melanization. A mutation in the N-terminus of processed CS in *P. berghei* changing Glutamine to Alanine and preventing the formation of pyro-Glutamic acid (Kolli et al., 2022), or mutations in CS pexel I/II domains in *P. yoelii* (Zhu et al., 2022) triggered the melanization of a significant number of mature oocysts, yet in both cases significant numbers of mutant oocysts still evaded melanization, suggesting that other parasite factors may also be conferring immune evasion. It is worth noting that these assays were done in *An. stephensi* and host immune responses may manifest differently in different *Plasmodium*-*Anopheles* combinations (Simoes et al., 2017). It would be interesting to test whether the melanization of these mutant rodent parasites will be further enhanced or even achieve complete refratoriness in ds*CTL4* or ds*CLIPA2*/*A14* melanotic backgrounds. In conclusion, melanization is a very complex phenomenon that is most likely regulated by a combination of host and parasite-specific factors.

In summary, we show for the first time that mature oocysts of wildtype *P. berghei* and *P. falciparum* parasites are susceptible to the mosquito melanization response to varying degrees in *An. gambiae*. Since ds*CLIPA2*/*A14* mosquitoes showed a higher melanization prevalence against *P. falciparum* oocysts compared to ookinetes, and that CTL4^null^ mosquitoes were shown to potently melanize *P. falciparum* ookinetes, a transgenic approach that targets both stages holds promise in achieving better transmission blocking potential

## Ethics statement

Work in this study was performed according to the recommendations in the Guide for the Care and Use of Laboratory Animals of the National Institutes of Health (Bethesda, USA). The followed animal protocol was approved by the Institutional Animal Care and Use committee (IACUC) of the American University of Beirut (permit number 18-08-504). The IACUC works in compliance with the Public Health Service Policy on the Humane Care and Use of Laboratory Animals (USA), and adopts the Guide for the Care and Use of Laboratory Animals of the National Institutes of Health.

## Acknowledgments

We would like to thank Sara Hajj Youssef for *Anopheles gambiae* G3 strain rearing and the Kamal A. Shair Central Research Laboratory at the American University of Beirut (AUB) for providing free access to their equipment. We would like to thank the Johns Hopkins Malaria Research Institute parasite and insectary core facilities for mosquitoes and *Plasmodium falciparum* culture. This work has been supported by the National Institutes of Health NIAD grant R01AI140760 and AUB Research Board award numbers 104261 and 104391 to M.A.O, and NIH/NIAD grants R01AI158615 and 1RO1AI170692, and the Bloomberg Philanthropies to G.D.

## Author contributions

SZ: Analysis, Investigation, Writing-original draft; SJ: Analysis, Investigation; SAS: Analysis, Investigation, Writing-original draft. JN: Analysis, Investigation; GD: Conceptualization, Funding acquisition, Supervision, Writing-review and editing; MAO: Conceptualization, Funding acquisition, Supervision, Project administration, Writing-original draft, review and editing.

