## Supplementary Material for "Late sporogonic stages of *Plasmodium* parasites are susceptible to the melanization response in *Anopheles gambiae* mosquitoes"

[*]**Correspondence:**

Mike A. Osta

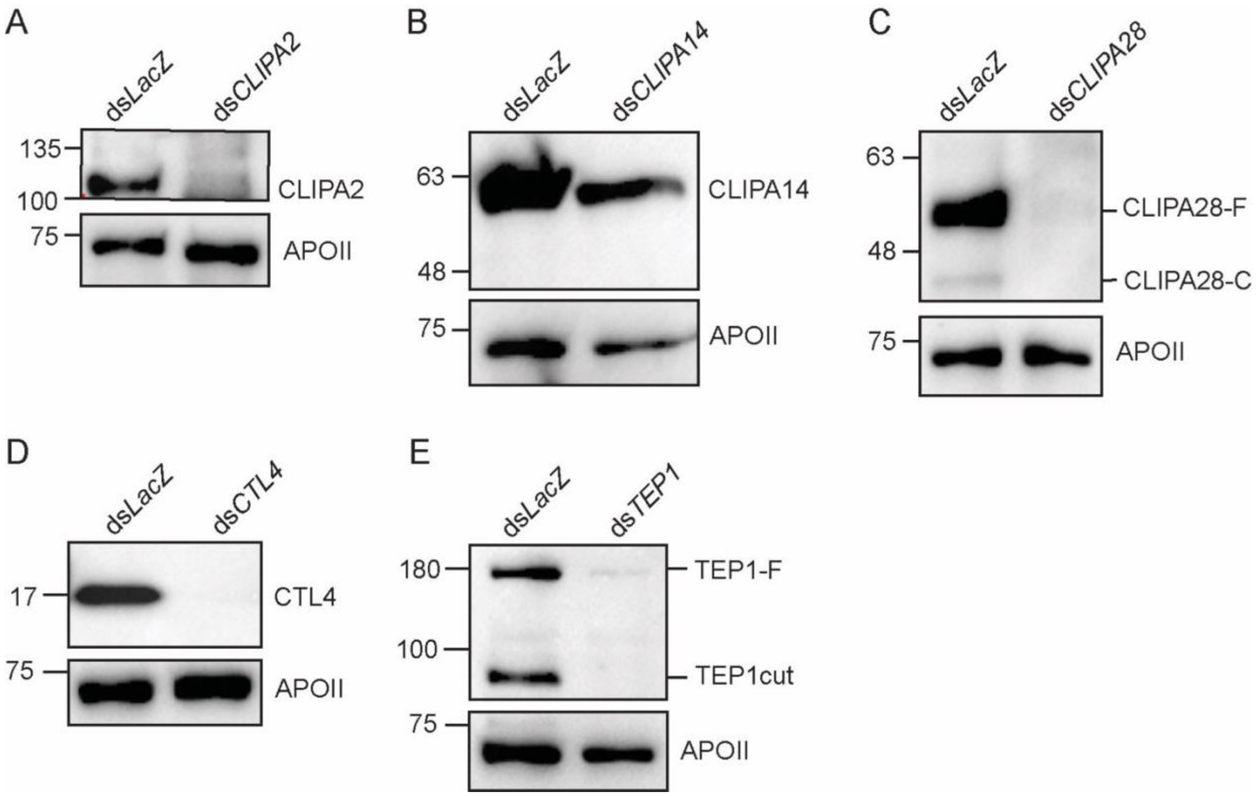


**Supplementary Figure 1. Knockdown efficiency of RNAi-silenced genes.** Shown are western blots of hemolymph samples extracted 7 days post-injection of mosquitoes with **(A)** ds*CLIPA2*, **(B)** ds*CLIPA14*, **(C)** ds*CLIPA28*, **(D)** ds*CTL4* and **(E)** ds*TEP1*. All hemolymph samples were extracted from 40 mosquitoes. Membranes were reprobed with αAPOII (without stripping) as loading control.

**Supplementary Table S1**

Raw data of Figure 1A: Independent experiments are shown in different colors

| | dsLacZ | | dsCLIPA2 | | dsCLIPA14 | | dsCLIPA2/A14 | |
|---|---|---|---|---|---|---|---|---|
| gut # | live oocysts | melanized oocysts | live oocysts | melanized oocysts | live oocysts | melanized oocysts | live oocysts | melanized oocysts |
| 1 | 3 | 0 | 7 | 1 | 3 | 0 | 43 | 22 |
| 2 | 33 | 0 | 8 | 0 | 3 | 1 | 14 | 1 |
| 3 | 29 | 0 | 7 | 0 | 1 | 0 | 16 | 17 |
| 4 | 237 | 0 | 0 | 0 | 0 | 0 | 95 | 141 |
| 5 | 14 | 0 | 1 | 0 | 114 | 0 | 9 | 0 |
| 6 | 10 | 0 | 54 | 0 | 2 | 0 | 1 | 0 |
| 7 | 39 | 0 | 34 | 1 | 267 | 1 | 0 | 1 |
| 8 | 81 | 0 | 5 | 0 | 4 | 3 | 89 | 13 |
| 9 | 5 | 1 | 2 | 0 | 32 | 0 | 113 | 8 |
| 10 | 1 | 0 | 5 | 0 | 247 | 2 | 0 | 8 |
| 11 | 12 | 0 | 7 | 0 | 2 | 0 | 12 | 8 |
| 12 | 6 | 0 | 4 | 0 | 0 | 0 | 6 | 2 |
| 13 | 78 | 0 | 0 | 0 | 1 | 2 | 96 | 57 |
| 14 | 30 | 0 | 246 | 0 | 21 | 1 | 66 | 54 |
| 15 | 52 | 0 | 5 | 0 | 52 | 2 | 10 | 0 |
| 16 | 210 | 0 | 15 | 0 | 22 | 2 | 10 | 0 |
| 17 | 327 | 0 | 6 | 0 | 65 | 4 | 149 | 45 |
| 18 | 514 | 0 | 167 | 0 | 8 | 0 | 43 | 101 |
| 19 | 325 | 0 | 1 | 0 | 32 | 0 | 150 | 10 |
| 20 | 40 | 0 | 14 | 0 | 13 | 2 | 78 | 57 |
| 21 | 436 | 0 | 53 | 0 | 126 | 16 | 0 | 0 |
| 22 | 0 | 0 | 0 | 0 | 12 | 6 | 81 | 31 |
| 23 | 14 | 0 | 149 | 0 | 41 | 0 | 288 | 16 |
| 24 | 5 | 0 | 215 | 1 | 1 | 0 | 67 | 59 |
| 25 | 15 | 0 | 34 | 0 | 23 | 7 | 46 | 1 |
| 26 | 0 | 0 | 0 | 0 | 25 | 9 | 118 | 40 |
| 27 | 15 | 0 | 36 | 0 | 0 | 0 | 9 | 20 |
| 28 | 64 | 0 | 0 | 0 | 442 | 0 | 12 | 21 |
| 29 | 262 | 0 | 0 | 0 | 87 | 30 | 0 | 0 |
| 30 | 77 | 0 | 4 | 0 | 26 | 0 | 0 | 0 |
| 31 | 13 | 0 | 0 | 0 | 2 | 0 | 17 | 0 |
| 32 | 0 | 1 | 0 | 0 | 12 | 0 | 0 | 4 |
| 33 | 0 | 1 | 137 | 0 | 34 | 0 | 21 | 30 |
| 34 | 31 | 0 | 1 | 0 | 14 | 0 | 0 | 0 |
| 35 | 0 | 0 | 8 | 0 | 62 | 5 | 8 | 5 |
| 36 | 5 | 0 | 0 | 0 | 2 | 0 | 0 | 0 |
| 37 | 27 | 0 | 0 | 0 | 2 | 0 | 0 | 1 |
| 38 | 10 | 0 | 7 | 0 | 4 | 0 | 2 | 0 |
| 39 | 3 | 0 | 0 | 0 | 30 | 10 | 0 | 0 |
| 40 | 4 | 0 | 0 | 0 | 0 | 0 | 0 | 2 |
| 41 | 48 | 0 | 0 | 0 | 0 | 1 | 15 | 24 |

| | | | | | | | | |
|---|---|---|---|---|---|---|---|---|
| 42 | 0 | 1 | 7 | 0 | 0 | 0 | 0 | 0 |
| 43 | 3 | 0 | 0 | 0 | 2 | 3 | 13 | 1 |
| 44 | 8 | 0 | 0 | 0 | 17 | 0 | 1 | 2 |
| 45 | 21 | 0 | 0 | 0 | 77 | 0 | 0 | 0 |
| 46 | 0 | 0 | 0 | 0 | 71 | 2 | 34 | 5 |
| 47 | 7 | 0 | 2 | 0 | 2 | 0 | 88 | 11 |
| 48 | 7 | 0 | | | 58 | 0 | 24 | 2 |
| 49 | 1 | 5 | | | 10 | 7 | 2 | 0 |
| 50 | 26 | 0 | | | 0 | 0 | 0 | 0 |
| 51 | 16 | 0 | | | 0 | 0 | 15 | 7 |
| 52 | 89 | 0 | | | 0 | 0 | 46 | 15 |
| 53 | 0 | 0 | | | 15 | 0 | 5 | 15 |
| 54 | 0 | 0 | | | | | 70 | 75 |
| 55 | 28 | 0 | | | | | 0 | 1 |
| 56 | 0 | 0 | | | | | 1 | 0 |
| 57 | 0 | 0 | | | | | 59 | 8 |
| 58 | 1 | 0 | | | | | 43 | 0 |
| 59 | 9 | 0 | | | | | 23 | 27 |
| 60 | 0 | 0 | | | | | 7 | 0 |
| 61 | 4 | 0 | | | | | | |
| 62 | 6 | 0 | | | | | | |
| 63 | 0 | 4 | | | | | | |

Raw data of Figure 1B: Independent experiments are shown in different colors

| | dsLacZ | | dsCLIPA2/A14 | | dsCTL4 | |
|---|---|---|---|---|---|---|
| Gut # | live oocysts | melanized oocysts | live oocysts | melanized oocysts | live oocysts | melanized oocysts |
| 1 | 25 | 0 | 16 | 0 | 1 | 0 |
| 2 | 20 | 2 | 14 | 0 | 3 | 0 |
| 3 | 28 | 0 | 4 | 1 | 134 | 0 |
| 4 | 114 | 3 | 19 | 33 | 19 | 0 |
| 5 | 5 | 0 | 16 | 2 | 15 | 0 |
| 6 | 93 | 0 | 19 | 50 | 3 | 0 |
| 7 | 4 | 0 | 5 | 0 | 30 | 1 |
| 8 | 39 | 0 | 54 | 0 | 86 | 0 |
| 9 | 7 | 0 | 9 | 3 | 1 | 0 |
| 10 | 2 | 0 | 1 | 1 | 2 | 0 |
| 11 | 10 | 0 | 17 | 1 | 23 | 0 |
| 12 | 4 | 0 | 8 | 0 | 37 | 0 |
| 13 | 158 | 0 | 6 | 3 | 0 | 0 |
| 14 | 15 | 3 | 0 | 0 | 7 | 0 |
| 15 | 61 | 0 | 13 | 12 | 1 | 0 |
| 16 | 49 | 0 | 4 | 1 | 1 | 0 |
| 17 | 42 | 0 | 1 | 0 | 5 | 0 |
| 18 | 34 | 0 | 3 | 10 | 0 | 0 |
| 19 | 1 | 0 | 6 | 28 | 0 | 0 |
| 20 | 28 | 1 | 0 | 0 | 5 | 0 |
| 21 | 144 | 0 | 153 | 8 | 0 | 0 |
| 22 | 4 | 0 | 8 | 0 | 34 | 0 |
| 23 | 0 | 0 | 88 | 16 | 10 | 0 |
| 24 | 4 | 0 | 1 | 1 | 37 | 2 |
| 25 | 14 | 2 | 1 | 2 | 3 | 0 |
| 26 | 3 | 0 | 0 | 0 | 12 | 0 |
| 27 | 24 | 0 | 14 | 11 | 4 | 0 |
| 28 | 145 | 0 | 69 | 0 | 150 | 0 |
| 29 | 36 | 0 | 85 | 1 | 56 | 0 |
| 30 | 4 | 0 | 30 | 34 | 0 | 0 |
| 31 | 0 | 0 | 6 | 2 | 9 | 0 |
| 32 | 2 | 0 | 53 | 11 | 54 | 1 |
| 33 | 67 | 0 | 12 | 13 | 11 | 5 |
| 34 | 4 | 0 | 43 | 62 | 6 | 0 |
| 35 | 1 | 0 | 14 | 20 | 4 | 0 |
| 36 | 72 | 0 | 32 | 0 | 87 | 0 |
| 37 | 12 | 0 | 25 | 65 | 33 | 0 |
| 38 | 14 | 0 | 23 | 3 | 8 | 0 |
| 39 | 13 | 0 | 5 | 14 | 17 | 0 |
| 40 | 21 | 0 | 6 | 14 | 240 | 0 |
| 41 | 15 | 0 | 60 | 14 | 0 | 2 |
| 42 | 2 | 0 | 0 | 2 | 31 | 0 |

| | | | | | | |
|---|---|---|---|---|---|---|
| 43 | 54 | 0 | 30 | 39 | 3 | 0 |
| 44 | 41 | 0 | 12 | 7 | 18 | 0 |
| 45 | 7 | 0 | 4 | 0 | 106 | 0 |
| 46 | 14 | 0 | 1 | 3 | 6 | 0 |
| 47 | 25 | 0 | 14 | 63 | 0 | 0 |
| 48 | 4 | 0 | 41 | 10 | 23 | 0 |
| 49 | 250 | 0 | 6 | 2 | 1 | 0 |
| 50 | 1 | 0 | 2 | 0 | 16 | 0 |
| 51 | 7 | 0 | 34 | 50 | 1 | 0 |
| 52 | 36 | 0 | 0 | 0 | 2 | 0 |
| 53 | 21 | 0 | 47 | 65 | 0 | 0 |
| 54 | 131 | 1 | 9 | 8 | 26 | 0 |
| 55 | 30 | 0 | 16 | 0 | 103 | 0 |
| 56 | 18 | 0 | 6 | 1 | 0 | 0 |
| 57 | 15 | 0 | 2 | 0 | 39 | 0 |
| 58 | 0 | 0 | 3 | 5 | 25 | 0 |
| 59 | 40 | 0 | 13 | 1 | 11 | 0 |
| 60 | 26 | 0 | 0 | 0 | 23 | 0 |
| 61 | 54 | 0 | 18 | 31 | 55 | 0 |
| 62 | 9 | 0 | 3 | 2 | | |
| 63 | 2 | 1 | 6 | 0 | | |
| 64 | 55 | 0 | 20 | 5 | | |
| 65 | 71 | 0 | 53 | 14 | | |
| 66 | 73 | 0 | 19 | 9 | | |
| 67 | 15 | 0 | 77 | 58 | | |
| 68 | 74 | 0 | 19 | 8 | | |
| 69 | 42 | 0 | 0 | 3 | | |
| 70 | 40 | 0 | 14 | 0 | | |
| 71 | 1 | 0 | 8 | 2 | | |
| 72 | 111 | 0 | 1 | 3 | | |
| 73 | 23 | 0 | 0 | 3 | | |
| 74 | 225 | 0 | 1 | 0 | | |
| 75 | 17 | 0 | 35 | 8 | | |
| 76 | 3 | 0 | 107 | 0 | | |
| 77 | | | 24 | 13 | | |
| 78 | | | 21 | 50 | | |
| 79 | | | 25 | 2 | | |
| 80 | | | 7 | 0 | | |
| 81 | | | 3 | 4 | | |

Raw data of Figure 1E:Independent experiments are shown in different colors

| | dsLacZ | | dsCLIPA2/A14 | |
|---|---|---|---|---|
| Gut # | live oocysts | melanized oocysts | live oocysts | melanized oocysts |
| 1 | 5 | 0 | 0 | 0 |
| 2 | 25 | 0 | 34 | 0 |
| 3 | 73 | 0 | 15 | 1 |
| 4 | 4 | 0 | 18 | 0 |
| 5 | 6 | 0 | 50 | 0 |
| 6 | 107 | 0 | 0 | 0 |
| 7 | 35 | 0 | 0 | 0 |
| 8 | 6 | 0 | 5 | 0 |
| 9 | 21 | 0 | 200 | 0 |
| 10 | 6 | 0 | 0 | 0 |
| 11 | 0 | 0 | 0 | 0 |
| 12 | 0 | 0 | 22 | 0 |
| 13 | 0 | 0 | 6 | 0 |
| 14 | 1 | 0 | 116 | 0 |
| 15 | 53 | 0 | 32 | 0 |
| 16 | 142 | 0 | 142 | 0 |
| 17 | 4 | 0 | 230 | 0 |
| 18 | 0 | 0 | 7 | 0 |
| 19 | 40 | 0 | 11 | 0 |
| 20 | 65 | 0 | 26 | 1 |
| 21 | 56 | 0 | 15 | 0 |
| 22 | 18 | 0 | 103 | 0 |
| 23 | 44 | 0 | 254 | 0 |
| 24 | 4 | 0 | 244 | 0 |
| 25 | 88 | 0 | 100 | 3 |
| 26 | 160 | 0 | 36 | 0 |
| 27 | 70 | 0 | 93 | 0 |
| 28 | 82 | 0 | 34 | 1 |
| 29 | 1 | 0 | 89 | 0 |
| 30 | 0 | 0 | 110 | 12 |
| 31 | 6 | 0 | 10 | 0 |
| 32 | 147 | 0 | 49 | 0 |
| 33 | 86 | 0 | 0 | 0 |
| 34 | 130 | 0 | 9 | 0 |
| 35 | 141 | 0 | 65 | 0 |
| 36 | 173 | 0 | 66 | 1 |
| 37 | 31 | 0 | 0 | 0 |
| 38 | 43 | 0 | 0 | 0 |
| 39 | 84 | 0 | 0 | 0 |
| 40 | 85 | 0 | 32 | 0 |
| 41 | 132 | 0 | 3 | 0 |
| 42 | 260 | 0 | 11 | 0 |
| 43 | 33 | 0 | 162 | 0 |

| | | | | |
|---|---|---|---|---|
| 44 | 1 | 0 | 158 | 0 |
| 45 | 19 | 0 | 28 | 0 |
| 46 | 0 | 13 | 0 | 2 |
| 47 | 152 | 0 | 2 | 0 |
| 48 | 22 | 0 | 3 | 0 |
| 49 | 0 | 0 | 17 | 3 |
| 50 | 12 | 0 | 0 | 0 |
| 51 | 0 | 0 | 0 | 0 |
| 52 | 30 | 0 | 15 | 0 |
| 53 | 0 | 0 | 0 | 0 |
| 54 | 0 | 0 | 124 | 0 |
| 55 | 142 | 0 | 1 | 0 |
| 56 | 1 | 0 | 0 | 0 |
| 57 | 151 | 0 | 0 | 0 |
| 58 | 31 | 0 | 73 | 0 |
| 59 | 0 | 0 | 22 | 0 |
| 60 | 83 | 0 | 96 | 0 |
| 61 | 186 | 0 | 0 | 0 |
| 62 | 29 | 0 | | |
| 63 | 0 | 0 | | |
| 64 | 0 | 0 | | |
| 65 | 31 | 0 | | |
| 66 | 0 | 0 | | |
| 67 | 0 | 1 | | |
| 68 | 40 | 0 | | |
| 69 | 33 | 0 | | |
| 70 | 3 | 0 | | |
| 71 | 0 | 0 | | |
| 72 | 0 | 0 | | |
| 73 | 0 | 0 | | |
| 74 | 46 | 0 | | |
| 75 | 0 | 0 | | |
| 76 | 0 | 0 | | |
| 77 | 2 | 0 | | |
| 78 | 1 | 0 | | |
| 79 | 13 | 0 | | |

Raw data of Figure 3: Independent experiments are shown in different colors

| | Sugar fed | | | | Extra blood meal | | | |
|---|---|---|---|---|---|---|---|---|
| | dsLacZ | | dsCLIPA2/A14 | | dsLacZ | | dsCLIPA2/A14 | |
| Gut # | live oocysts | melanized oocysts | live oocysts | melanized oocysts | live oocysts | melanized oocysts | live oocysts | melanized oocysts |
| 1 | 10 | 0 | 53 | 1 | 0 | 0 | 24 | 47 |
| 2 | 42 | 2 | 1 | 0 | 147 | 0 | 23 | 7 |
| 3 | 11 | 0 | 33 | 5 | 0 | 0 | 1 | 0 |
| 4 | 81 | 0 | 0 | 0 | 0 | 0 | 2 | 2 |
| 5 | 40 | 0 | 2 | 1 | 1 | 0 | 0 | 0 |
| 6 | 0 | 0 | 5 | 7 | 12 | 11 | 14 | 18 |
| 7 | 2 | 0 | 42 | 0 | 84 | 1 | 15 | 4 |
| 8 | 13 | 0 | 5 | 8 | 7 | 0 | 43 | 112 |
| 9 | 200 | 0 | 12 | 31 | 37 | 3 | 8 | 9 |
| 10 | 35 | 0 | 16 | 0 | 11 | 0 | 14 | 24 |
| 11 | 2 | 0 | 2 | 0 | 8 | 0 | 5 | 0 |
| 12 | 80 | 0 | 101 | 5 | 6 | 0 | 203 | 23 |
| 13 | 180 | 0 | 22 | 0 | 0 | 0 | 0 | 0 |
| 14 | 210 | 0 | 15 | 15 | 2 | 0 | 0 | 0 |
| 15 | 0 | 0 | 134 | 18 | 37 | 2 | 43 | 4 |
| 16 | 11 | 0 | 0 | 0 | 15 | 0 | 2 | 0 |
| 17 | 1 | 0 | 1 | 1 | 90 | 1 | 23 | 7 |
| 18 | 94 | 0 | 1 | 5 | 0 | 0 | 2 | 0 |
| 19 | 80 | 1 | 4 | 0 | 20 | 0 | 2 | 0 |
| 20 | 73 | 0 | 14 | 27 | 8 | 0 | 151 | 19 |
| 21 | 67 | 0 | 71 | 15 | 26 | 0 | 85 | 74 |
| 22 | 27 | 0 | 62 | 22 | 0 | 0 | 3 | 0 |
| 23 | 14 | 0 | 30 | 6 | 8 | 0 | 2 | 0 |
| 24 | 0 | 0 | 22 | 29 | 8 | 0 | 1 | 13 |
| 25 | 3 | 0 | 250 | 30 | 11 | 0 | 17 | 10 |
| 26 | 73 | 0 | 49 | 22 | 2 | 0 | 52 | 30 |
| 27 | 44 | 0 | 15 | 11 | 1 | 1 | 26 | 28 |
| 28 | 1 | 0 | 14 | 19 | 12 | 0 | 21 | 28 |
| 29 | 16 | 0 | 17 | 6 | 8 | 1 | 0 | 0 |
| 30 | 19 | 0 | 2 | 0 | 0 | 0 | 64 | 139 |
| 31 | 3 | 0 | 0 | 1 | 0 | 0 | | |
| 32 | 6 | 0 | | | 67 | 0 | 30 | 96 |
| 33 | 12 | 0 | 3 | 1 | 0 | 0 | 15 | 20 |
| 34 | 20 | 0 | 62 | 39 | 11 | 0 | 6 | 4 |
| 35 | 4 | 0 | 9 | 18 | | | 31 | 32 |
| 36 | 39 | 0 | 46 | 43 | | | 41 | 20 |
| 37 | 10 | 0 | 116 | 50 | | | 54 | 32 |
| 38 | 23 | 0 | 5 | 6 | | | 2 | 2 |
| 39 | 5 | 0 | 18 | 37 | | | 61 | 62 |
| 40 | 0 | 0 | 40 | 10 | | | 0 | 0 |
| 41 | 67 | 0 | 20 | 19 | | | 2 | 2 |

| | | | | | | |
|---|---|---|---|---|---|---|
| 42 | 80 | 0 | 10 | 0 | 72 | 42 |
| 43 | 160 | 0 | 126 | 28 | 0 | 0 |
| 44 | 23 | 0 | 7 | 61 | 32 | 52 |
| 45 | 15 | 0 | 1 | 1 | 22 | 2 |
| 46 | 19 | 0 | 5 | 18 | 9 | 54 |
| 47 | 35 | 0 | 85 | 69 | 1 | 10 |
| 48 | 14 | 0 | 21 | 12 | 13 | 21 |
| 49 | | | 0 | 0 | 0 | 0 |
| 50 | | | 14 | 1 | 2 | 6 |
| 51 | | | | | 18 | 0 |
| 52 | | | | | 77 | 74 |
| 53 | | | | | 51 | 75 |
| 54 | | | | | 117 | 65 |
| 55 | | | | | 38 | 96 |
| 56 | | | | | 3 | 27 |
| 57 | | | | | 6 | 12 |
| 58 | | | | | 16 | 34 |
| 59 | | | | | 0 | 0 |
| 60 | | | | | 8 | 5 |
| 61 | | | | | 74 | 128 |
| 62 | | | | | 4 | 20 |
| 63 | | | | | 1 | 5 |
| 64 | | | | | 22 | 29 |
| 65 | | | | | 16 | 47 |
| 66 | | | | | 18 | 15 |
| 67 | | | | | 43 | 44 |

Raw data of figure 4A:Independent experiments are shown in different colors

| | dsLacZ | | dsCLIPA2/A14 | | dsCLIPA2/A14/A28 | | dsCLIPA2/A14/TEP1 | |
|---|---|---|---|---|---|---|---|---|
| Gut # | live oocysts | melanized oocysts | live oocysts | melanized oocysts | live oocysts | melanized oocysts | live oocysts | melanized oocysts |
| 1 | 68 | 0 | 9 | 15 | 26 | 0 | 104 | 0 |
| 2 | 58 | 1 | 13 | 7 | 284 | 0 | 2 | 0 |
| 3 | 140 | 0 | 10 | 0 | 111 | 0 | 1 | 0 |
| 4 | 339 | 0 | 3 | 1 | 100 | 0 | 17 | 0 |
| 5 | 16 | 0 | 113 | 5 | 125 | 0 | 67 | 0 |
| 6 | 14 | 0 | 39 | 77 | 144 | 3 | 1 | 0 |
| 7 | 82 | 0 | 17 | 34 | 38 | 0 | 4 | 0 |
| 8 | 85 | 2 | 1 | 0 | 29 | 0 | 0 | 2 |
| 9 | 89 | 0 | 4 | 0 | 0 | 0 | 102 | 7 |
| 10 | 3 | 0 | 92 | 20 | 0 | 0 | 8 | 0 |
| 11 | 133 | 0 | 50 | 36 | 52 | 0 | 77 | 0 |
| 12 | 1 | 0 | 35 | 22 | 54 | 0 | 48 | 0 |
| 13 | 26 | 0 | 0 | 0 | 163 | 0 | 4 | 14 |
| 14 | 13 | 0 | 8 | 0 | 66 | 0 | 0 | 0 |
| 15 | 65 | 0 | 10 | 20 | 140 | 0 | 36 | 0 |
| 16 | 121 | 0 | 7 | 8 | 24 | 0 | 19 | 0 |
| 17 | 27 | 0 | 57 | 30 | 106 | 0 | 108 | 1 |
| 18 | 0 | 0 | 26 | 1 | 4 | 0 | 0 | 0 |
| 19 | 39 | 0 | 0 | 0 | 42 | 1 | 7 | 0 |
| 20 | 2 | 0 | 13 | 0 | 6 | 0 | 4 | 1 |
| 21 | 1 | 0 | 13 | 14 | 4 | 2 | 35 | 1 |
| 22 | 32 | 0 | 38 | 80 | 34 | 0 | 7 | 1 |
| 23 | 3 | 3 | 8 | 21 | 1 | 2 | 13 | 0 |
| 24 | 102 | 0 | 6 | 25 | 5 | 0 | 11 | 0 |
| 25 | 36 | 0 | 26 | 6 | 1 | 0 | 72 | 0 |
| 26 | 1 | 0 | 7 | 0 | 1 | 1 | 9 | 1 |
| 27 | 0 | 0 | 11 | 11 | 16 | 0 | 29 | 0 |
| 28 | 14 | 0 | 9 | 1 | 8 | 0 | 1 | 0 |
| 29 | 3 | 0 | 2 | 16 | 2 | 0 | 4 | 0 |
| 30 | 24 | 1 | 3 | 3 | 5 | 0 | 108 | 0 |
| 31 | 35 | 0 | 18 | 29 | 34 | 0 | 2 | 0 |
| 32 | 64 | 0 | 0 | 0 | 82 | 0 | 2 | 0 |
| 33 | 6 | 0 | 2 | 21 | 35 | 1 | 22 | 1 |
| 34 | 15 | 0 | 0 | 0 | 15 | 0 | 2 | 0 |
| 35 | 16 | 0 | 33 | 50 | 89 | 0 | 3 | 0 |
| 36 | 4 | 0 | 32 | 3 | 97 | 1 | 25 | 1 |
| 37 | 24 | 0 | 1 | 5 | 87 | 0 | 5 | 2 |
| 38 | 3 | 0 | 32 | 30 | 115 | 0 | 0 | 0 |
| 39 | 0 | 0 | 29 | 33 | 11 | 0 | 8 | 0 |
| 40 | 6 | 0 | 2 | 8 | 1 | 0 | 46 | 0 |
| 41 | | | | | 39 | 0 | 12 | 0 |
| 42 | | | | | 19 | 0 | 10 | 0 |

| | | | | | |
|---|---|---|---|---|---|
| 43 | | 10 | 0 | 3 | 0 |
| 44 | | 0 | 0 | 12 | 0 |
| 45 | | 4 | 0 | 5 | 1 |
| 46 | | 48 | 0 | 5 | 0 |
| 47 | | 31 | 0 | 42 | 0 |

**Supplementary Table 2. Primers used for dsRNA production**

| Gene | Primer sequence (T7 promoter sequence underlined) | Reference |
|---|---|---|
| LacZ | For: 5'-TAATACGACTCACTATAGGGAGAATCCGACGGGTTGTTACT-3'<br>Rev: 5'-TAATACGACTCACTATAGGGCACCACGCTCATCGATAATTT-3' | (Habtewold et al., 2008) |
| GFP | For: 5'- TAATACGACTCACTATAGGGTTCATCTGCACCACCGGC-3'<br>Rev: 5'-TAATACGACTCACTATAGGGCTGGTAGTGGTCGGCGAG-3' | (Simoes et al., 2017) |
| CLIPA2<br>(AGAP011790) | For:5'-TAATACGACTCACTATAGGGATCCTAACAACGGCACACTGTGTGA-3'<br>Rev:5'-TAATACGACTCACTATAGGGTCCTGATCGCCATGATTGGTGGTGCT-3' | (Yassine et al., 2014) |
| CLIPA14<br>(AGAP011788) | For: 5'- TAATACGACTCACTATAGGGCGGCATCATCGACATCCGTGTC-3'<br>Rev: 5'- TAATACGACTCACTATAGGGGTTGCTGTCGGCGACACGCTCCT-3' | (Nakhleh et al., 2017) |
| CLIPA28<br>(AGAP010730) | For: 5'-TAATACGACTCACTATAGGGAGACCACCAAGGAACCGTTCCCGCAGCAA-3'<br>Rev: 5'-TAATACGACTCACTATAGGGAGACCGCAACCGATGCCCCACGATACGAT-3' | (El Moussawi et al., 2019) |
| TEP1<br>(AGAP010815) | For: 5'-TAATACGACTCACTATAGGGTTTGTGGGCCTTAAAGCGCTG-3'<br>Rev: 5'- TAATACGACTCACTATAGGGACCACGTAACCGCTCGGTAAG-3' | (Povelones et al., 2011) |
| CTL4<br>(AGAP005335) | For: 5'- TAATACGACTCACTATAGGGGTTAGCAGCATTGGGATTACCCT-3'<br>Rev: 5'- TAATACGACTCACTATAGGGGAAGTCGCAACCCAGCTCATTGT-3' | (Povelones et al., 2013) |

| **Supplementary Table 3. Infection data for *P. falciparum* experiments** | | | | | |
|---|---|---|---|---|---|
| Experiment | Silenced gene | parasite | N | Median L, M | Prevalence % |
| Early-stage melanization (Low median) | GFP | Pf NF54 | 127 | 6, 0 | 79 |
| | A2/A14 | Pf NF54 | 117 | 9, 0 | 79 |
| Early-stage melanization (High median) | GFP | Pf NF54 | 63 | 64,0 | 94 |
| | A2/A14 | Pf NF54 | 61 | 68,0 | 97 |
| Late stage melanization | GFP | Pf NF54 | 167 | 7,0 | 84 |
| | A2/A14 | Pf NF54 | 143 | 4,0 | 77 |
